# Role of mesolimbic cannabinoid receptor 1 in stress-driven increases in cocaine self-administration in male rats

**DOI:** 10.1101/2022.10.28.514315

**Authors:** Jayme R. McReynolds, Colten P. Wolf, Dylan M. Starck, Jacob C. Mathy, Rebecca Schaps, Leslie A. Krause, Cecilia J. Hillard, John R. Mantsch

**Author notes:** <u>Corresponding Author:</u> Jayme R. McReynolds, Address: 560 N 16^th^ St SC446, Milwaukee, WI 53233. Jayme R. McReynolds, Address: 2120 E. Galbraith Rd #A-141, Cincinnati, OH 45237.

## Abstract

Stress is prevalent in the lives of those with substance use disorders (SUDs) and influences SUD outcomes. Understanding the neurobiological mechanisms through which stress promotes drug use is important for the development of effective SUD interventions. We have developed a model wherein exposure to a stressor, uncontrollable electric footshock, daily at the time of cocaine self-administration (SA) escalates intake in male rats. Here we test the hypothesis that stress-induced escalation of cocaine SA requires CB1 cannabinoid receptor signaling. Male Sprague-Dawley rats self-administered cocaine (0.5 mg/kg/inf, i.v.) during 2-h sessions comprised of four 30-min SA components separated by 5-min shock sequences or 5-min shock-free periods for 14 days. Footshock produced an escalation of cocaine SA that persisted following shock removal. Systemic administration of the cannabinoid receptor type 1 (CB1R) antagonist, AM251, attenuated cocaine intake only in rats with a history of stress. This effect was localized to the mesolimbic system, as intra-nucleus accumbens (NAc) shell and intra-ventral tegmental area (VTA) micro-infusions of AM251 attenuated cocaine intake only in stress-escalated rats. Cocaine SA, regardless of stress history, increases CB1R binding site density in the VTA, but not NAc shell. Following extinction, cocaine-primed reinstatement (10 mg/kg, ip) was increased in rats with prior footshock during SA. AM251 attenuated reinstatement only in rats with a stress history. Altogether, these data demonstrate that mesolimbic CB1R signaling is required to escalate intake and heighten relapse susceptibility and suggest that repeated stress at the time of cocaine use enhances mesolimbic CB1R signaling through a currently unknown mechanism.

## Introduction

Despite intensive research efforts, substance use disorders (SUDs) remain among society’s most significant public health challenges, and there is a pressing need for more effective interventions. While emphasis has recently been placed on opioids, cocaine use is a persistent problem. There is a strong link between stress and cocaine use. Co-morbidity between cocaine use disorder (CUD) and stress-related conditions [1], notably PTSD, is high [2]. Measures of cumulative lifetime stress positively correlate with CUD severity [3]. Life stress often precedes cocaine use [4-6] and can predict relapse [7; 8] and poor treatment outcomes [9]. Further, relapse susceptibility is greater in those with lower distress tolerance [10]. In a lab setting, personalized stress imagery can precipitate cocaine craving [11; 12] and the magnitude of craving and associated physiological responses predicts subsequent relapse [8; 13].

A key characteristic of SUDs is a progressive loss of control over drug intake which can be observed in rodents as escalating patterns of drug self-administration (SA) [14; 15]. We have demonstrated that delivery of electric footshock stress, daily at the time of SA, produces an emergent escalation of cocaine intake in male rats [16]. Shock-induced escalation of cocaine SA involves neuroadaptations that require elevated glucocorticoids [16], as does escalation of SA in rats provided daily long-access to the drug [17; 18]. Endocannabinoid signaling is glucocorticoid-regulated [19-24], is involved in buffering stress responses and is affected by chronic stress [25; 26]. Polymorphisms in the cannabinoid type 1 receptor (CB1R) gene, *cnr1*, are associated with CUD [27]. In rodents, CB1R antagonism has minimal effects on cocaine SA under conditions of stable/non-escalated intake [28-31]. However, in rats displaying escalated intake following repeated daily long-access SA, the CB1R antagonist, rimonabant, reduced breakpoints when tested for cocaine SA under a progressive ratio schedule that are not observed after daily short-access SA [31]. Altogether these findings suggest that endocannabinoid signaling via CB1Rs is regulated by stress, recruited under conditions resulting in escalated drug use, and promotes drug SA. Accordingly, we hypothesize that stressor-induced escalation of cocaine SA involves heightened CB1R signaling in brain regions implicated in cocaine use.

VTA (ventral tegmental area) dopamine neurons projecting to the nucleus accumbens (NAc) shell comprise a mesolimbic pathway that is critical for cocaine SA [32-35]. Dopaminergic signaling is regulated in both regions by endocannabinoids [36-41]; and both cocaine and stress can influence VTA and NAc endocannabinoid signaling [31; 42-44]. Indeed, escalated cocaine intake observed under long-access SA conditions is associated with an upregulation of NAc shell CB1Rs and is selectively reduced by intra-NAc shell micro-infusions of rimonabant [31]. Here we test the hypothesis that repeated stress at the time of cocaine SA requires VTA and/or NAc shell CB1R signaling to escalate cocaine use. Further, we hypothesize that this involvement of CB1R signaling is long-lasting and heightens susceptibility to later cocaine-primed reinstatement.

## Materials and Methods

Additional methodological details are described in the supplemental methods.

### Subjects

155 Male Sprague-Dawley rats (60 days old; 275-300g at arrival; Envigo, Indianapolis, IN) were housed individually in a temperature- and humidity-controlled AAALAC-accredited animal facility on a 12:12 hr reverse light/dark cycle (0700-1900 lights off) with *ad libitum* food and water access. Procedures were conducted in the dark/active phase of the rat light cycle. Procedures were approved by the Institutional Animal Care and Use Committee at Marquette University and conducted in compliance with NIH Guidelines.

### Surgery

All rats received intravenous catheters for drug SA. Some rats also received intra-NAc shell or intra-VTA guide cannula immediately following the catheter surgery. Guide cannula were implanted 0.5 mm above the target brain region. Rats recovered for a minimum of one week before the commencement of SA training.

### Cocaine SA

Rats were trained to self-administer cocaine (0.5 mg/kg/0.2 ml infusion over 5 sec) by lever pressing under a fixed ratio 4 schedule during daily sessions as described [16] (Fig 1A). The program consisted of four 30-min SA blocks wherein the levers were extended, the active lever light was on, and the houselight was off. During the infusion, the active lever light was extinguished for a 10-sec time-out period during which presses were recorded but not reinforced. Separating the four 30-min SA blocks (and preceding the first block) were 5-min drug-free periods during which the houselight was on, the levers were retracted, and the active lever light was off.

**Figure 1.**
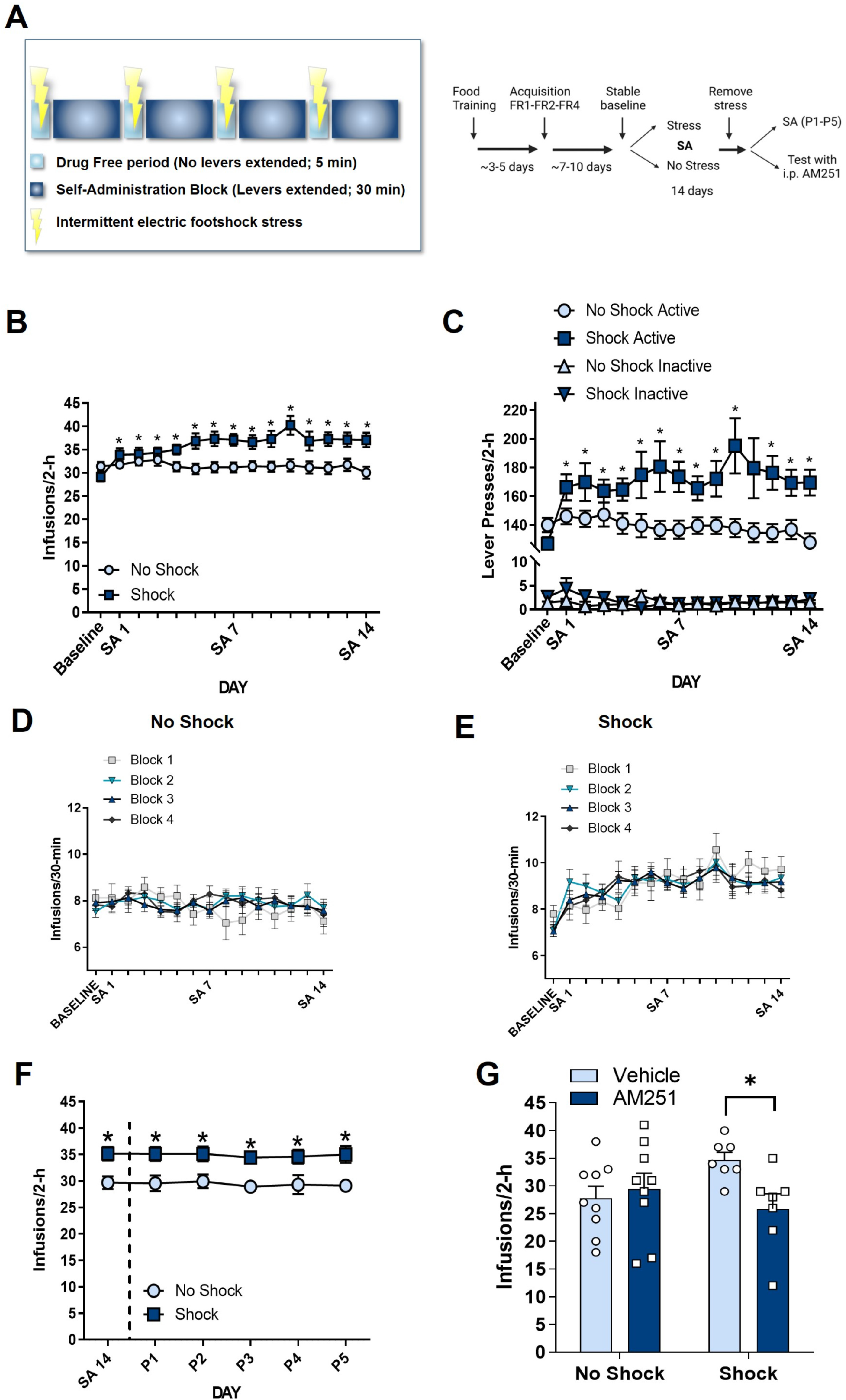
Chronic electronic footshock administered daily at the time of self-administration induces a persistent escalation of cocaine intake. A) Schematic of modified short-access paradigm and experimental timeline wherein 2-hr daily self-administration was separated into 4 30-min self-administration blocks (levers extended, drug cue light on, house light off) separated by 4 5-min drug-free periods (levers retracted, drug cue light off, house light on). Following stable self-administration on an FR4 schedule of reinforcement, animals in the stress group received intermittent electric footshock stress during the drug-free period daily for 14 days. Under this paradigm footshock (shock) stress administered daily at the time of self-administration induced a significant escalation of cocaine intake as assessed by an increase in infusions (B) and active lever presses (C) across days while no shock rats showed fairly stable responding. Daily infusions of cocaine were similar across all 4 self-administration blocks within each day in both the no shock (D) and shock (E) groups (no shock: n=20; shock n=17; *p<.05, compared to baseline). F) Animals in the stress group continue to show elevated cocaine intake when the shock stressor was removed (P1-P5; No Shock, n=13; Shock, n= 15; *p<.05, compared to No Shock). G) Systemic cannabinoid receptor type 1 (CB1R) antagonism with AM251 (0, 1 mg/kg, i.p.) given 30 min prior to self-administration significantly attenuated cocaine intake in animals with a history of shock (n=7) while having no effect in rats with a history of no shock (n=9; *p<.05 compared to Vehicle). Data are presented as mean ± SEM.

### Experiment 1: Electric footshock stress-induced escalation of cocaine intake

Rats were separated into two groups following establishment of baseline SA (<10% change in infusions over 3 days) (Fig. 1A). The “shock” group received intermittent electric footshock stress (3 × 0.4 mA, 500 ms duration, 30-s average inter-shock interval) during the four 5-min drug-free components over the 14-day SA period. The “no shock” group continued to self-administer cocaine for 14 days under the conditions described above where on each SA day, they had interleaving drug-free periods between 30-min SA blocks where the levers were retracted for the duration of the 5-min period and no footshock was administered. To test whether stress-induced escalation of cocaine intake persisted in the absence of the stressor, the shock component was removed after the 14-day SA period and the rats were allowed to self-administer cocaine for at least 5 days under “no shock” conditions. Control rats continued to self-administer under the same parameters during this time.

### Experiment 2: Effect of systemic CB1R antagonism on stress-escalated cocaine intake

Rats self-administered cocaine for 14 days after which the shock components were removed for at least one day prior to testing with AM251, a CB1R antagonist/inverse agonist. During testing, rats received injections of AM251 (1 mg/kg, i.p.) or vehicle (1:1:18 ethanol: Cremaphor: saline) 30 min before the SA session (Fig. 1G). All rats received both the vehicle and AM251 on different days in a within-subjects design in a counter-balanced order.

### Experiment 3: Effect of CB1R antagonism in the NAc shell and VTA on stress-escalated cocaine intake

Following 14 days of SA under no shock/shock conditions, rats received bilateral intra-NAc shell or intra-VTA AM251 (1, 3 µg/0.3 µL/side) or vehicle (100% DMSO) micro-infusions were tested for SA under shock-free conditions (15-min pretreatment; see Experiment 2). Each rat received both AM251 and vehicle in a within-subjects design in a counter-balanced order with at least one cocaine SA day between drug treatments to ensure normal responding before the next test. After the experiment, rats were euthanized, and brains were collected for cannula placement verification. Only rats with confirmed placements were included for analyses.

### Experiment 4: CB1R binding changes in the NAc shell and VTA following stress-induced escalation of cocaine intake

To examine CB1R binding, a binding assay was performed in postmortem tissue. Rats self-administered under no shock/shock conditions (see Experiment 1). Saline control groups were added to isolate changes in binding resulting from cocaine SA or repeated shock alone. Saline access was provided under parameters otherwise identical to those for cocaine. Rats were euthanized, brains were collected and frozen 24h after the last session. NAc shell and VTA were isolated from frozen brains and a 6-point homogenate CB1R binding assay in membrane fractions using [^3^H]CP55,940, a CB1R agonist, was performed (Supplemental Methods).

### Experiment 5: Cocaine-primed reinstatement following stress-induced escalation of cocaine intake; requirement for CB1R

Following 14 days of SA, rats underwent extinction training wherein cocaine was replaced with saline and the footshock stress was no longer administered. Once extinction criteria were met (<15 lever presses/2h), reinstatement testing was conducted during which rats received cocaine injections (0, 2.5, 5, 10 mg/kg, i.p.) immediately prior to the test session. Each rat received all doses in counter-balanced sequence. Rats met extinction criteria between consecutive reinstatement tests. To test the role of CB1R signaling in enhanced cocaine-induced reinstatement, additional groups of rats underwent SA and extinction/reinstatement. On reinstatement test day, rats received AM251 (1 mg/kg, i.p.) or vehicle followed by a cocaine (10 mg/kg, i.p.) or saline injection 30 min later, immediately followed by reinstatement testing. The sequence of AM251/vehicle tests was counter-balanced.

### Statistical analysis

Statistical analyses were conducted using 2-way or 3-way repeated measures ANOVA, followed by Dunnett’s or Holm-Sidak-corrected post hoc tests when appropriate.

## Results

For all experiments, relevant statistical analyses are reported in the main text and full statistical analyses are reported in supplemental tables (Tables S1-4).

### Experiment 1: Electric footshock stress-induced escalation of cocaine intake

Study design is shown in Fig 1a. Rats receiving footshock during cocaine SA, but not no-shock controls, showed significant escalation of cocaine SA (Fig. 1b-e). The cocaine self-administration data associated with Fig. 1F, Fig. 1G, and Fig.4 are represented in Fig. 1b-e. Two-way ANOVA revealed a significant stress x SA day interaction for cocaine infusions (Fig. 1B; F_(14,700)_=4.719, p<.0001) and active (Fig. 1C; F_(14, 700)_=2.561, p<.01), but not inactive, lever presses (Fig. 1C; F_(14, 700)_=1.539, p=.09). There were no significant differences in baseline SA between the groups; and shocked but not control rats showed significantly increased intake (p<.01) and active lever pressing (p<.01) on each SA day compared to baseline. Additionally, rats in the shock group had significantly greater intake relative to controls on SA days 5-14 (p<.05) as well as greater cumulative cocaine intake (Fig. S1A; t_(50)_=3.18, p<.01) and area under the curve (AUC; Fig. S1B; t_(50)_=5.14, p<.0001). Intake across the four SA blocks each day was consistent in both groups (Fig. 1D-E); there were no SA day x block interactions (No shock: F_(42,966)_=1.14, p=0.25; Shock: F_(42,1134)_=1.34, p=.07). To determine whether escalated intake persists in the absence of shock, rats in the shock group were allowed to continue SA under shock-free conditions for at least 5 days (Fig. 1F). These rats displayed escalated SA during this persistence phase (P1-5), comparable to when shock was present (SA 14). Two-way ANOVA revealed no significant SA day x stress interaction (F_(5,130)_=0.22, p=0.95) but an overall stress effect (F_(1,26)_=7.43, p<.05).

### Experiment 2: Effect of systemic CB1R antagonism on stress-escalated cocaine intake

To examine CB1R involvement in stress-escalated SA, the effects of AM251 were determined. Rats received AM251 (1 mg/kg, i.p.) or vehicle injections 30 min prior to SA. Two-way ANOVA revealed a significant AM251 x stress interaction (F_(1,14)_=18.56, p<.001); AM251 significantly attenuated intake in the shock (p<.001) but not control group.

### Experiment 3: Effect of CB1R antagonism in the NAc shell and VTA on stress-escalated cocaine intake

In this study, rats received micro-infusions of AM251 into NAc shell or VTA following 14 days of cocaine SA under stress or non-stress conditions (Fig 2A). As expected, SA was established in both groups and two-way ANOVA revealed a significant overall effect of stress (F_(1,33)_=12.09, p<.0001; Fig 2B). Rats in the shock group had significantly greater cumulative cocaine intake (Fig. S2C; t_(33)_=3.58, p<.01) and AUC (Fig. S2D; t_(33)_=2.47, p<.05) compared to controls. To test for involvement of NAc shell CB1R signaling, rats received intra-NAc shell micro-infusions of AM251 (1, 3 µg/side) or vehicle 15 min before SA (Fig. 2C). Three-way ANOVA revealed a significant drug x stress x dose interaction (F_(1,28)_=7.59, p<.05). AM251 did not attenuate SA in non-shocked controls at either dose (p>.05). However, both AM251 doses attenuated SA in shocked rats (1 µg: p<.01; 3 µg: p<.01). Separate rats received intra-VTA AM251 micro-infusions (1, 3 µg/side) or vehicle (Fig. 2E). While there was no effect of AM251 dose, there was an overall AM251 effect (F_(1,23)_=22.11, p<.001) and a significant AM251 x stress interaction (F_(2,23)_=4.31, p<.05). High dose AM251 (3 µg) attenuated SA in controls (p<.05), but both AM251 doses attenuated SA in shocked rats (1 µg: p=.05; 3 µg: p<.05).

**Figure 2.**
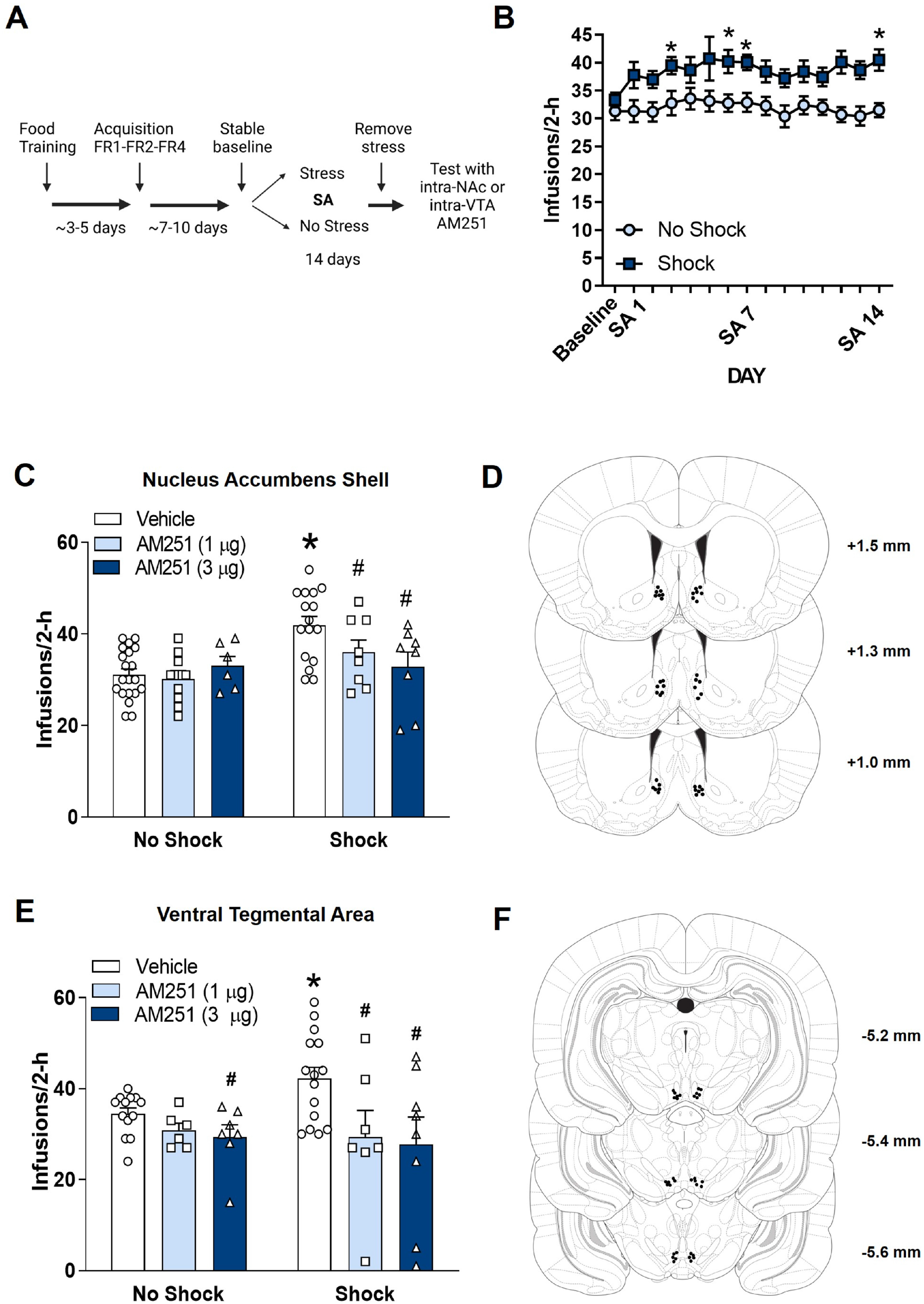
Cannabinoid receptor type 1 antagonism in the nucleus accumbens shell and ventral tegmental area attenuate cocaine intake in rats that underwent stress-induced escalation of cocaine intake. A) Experimental timeline. B) Self-administration data for all animals that received intra-cranial CB1R antagonism. Repeated daily footshock stress induces a significant escalation of cocaine intake (No shock, n=16; Shock, n=19; *p<.05 compared to baseline). C) Intra-nucleus accumbens (NAc) shell administration of the CB1R antagonist AM251 (0, 1, 3 µg/side) attenuates cocaine intake only in rats with a history of shock (n=8/dose) but has no effect on cocaine intake in rats with a history of no shock (n=7-9/dose; *p<.05 compared to No Shock-Vehicle, #p<.05 compared to Vehicle). D) Injection needle tips of all rats included in the intra-NAc shell experiment. Coordinates are listed from Bregma. E) Intra-ventral tegmental area (VTA) administration of AM251 (0, 1, 3 µg/side) attenuates cocaine intake in rats with a history of shock (n=7-8/dose) at both doses of AM251 but only attenuates cocaine intake in rats with a history of no shock (n=6-7/dose) at the 3-µg dose (*p<.05 compared to No Shock-Vehicle, #p<.05 compared to Vehicle). F) Injection needle tips of all rats included in the intra-VTA experiment. Coordinates are listed from Bregma. Data are presented as mean ± SEM.

### Experiment 4: CB1R binding changes in the NAc shell and VTA following stress-induced escalation of cocaine intake

CB1R binding parameters were determined in membranes harvested from VTA and NAc shell 24 hrs after the last SA day in shocked and control rats. Separate groups were included that had saline access for SA with or without shock. SA data from rats used for this study are shown in Fig. 3A. Two-way ANOVA with SA condition as a between-subject factor and day as a within-subject factor revealed a significant SA condition x day interaction (F_(42,700)_=1.58, p<.05). For cocaine SA, rats in the shock group had significantly greater cumulative cocaine intake (Fig. S3C; t_(25)_=2.56, p<.05) compared to controls. For AUC, a 2-way ANOVA revealed a significant SA condition x shock interaction (Fig. S3D; F_(1,45)_=4.52, p<.05); AUC in the cocaine SA/shock group was significantly increased compared to every other group (p<.05). In the NAc shell, all data sets (F_(6,263)_=1.38, p=0.22) were fit by the same curve, indicating that the binding parameters are not significantly different among the treatment groups (Fig. 3C). By contrast, in the VTA (Fig. 3E), one curve could not adequately fit all data sets (F_(6,233)_=2.93, p<.01), and B_max_ and K_D_ values were determined from the mean binding data for each group (Table S5). K_D_ values did not differ across groups, and there was high overall variance in the calculated B_max_ values, likely because binding site density in the VTA is very low. As an additional approach, we compared specific binding at the highest ^3^H-CP 55,940 concentration in each sample using two-way ANOVA. There was a significant overall effect of SA condition (F_(1,41)_=74.49, p<.01), with any history of cocaine SA producing higher amounts of ^3^H-CP55940 binding (Fig. 3F).The individual binding values at the highest ^3^H-CP 55,940 concentration in the NAc shell are shown in Fig. 3D.

**Figure 3.**
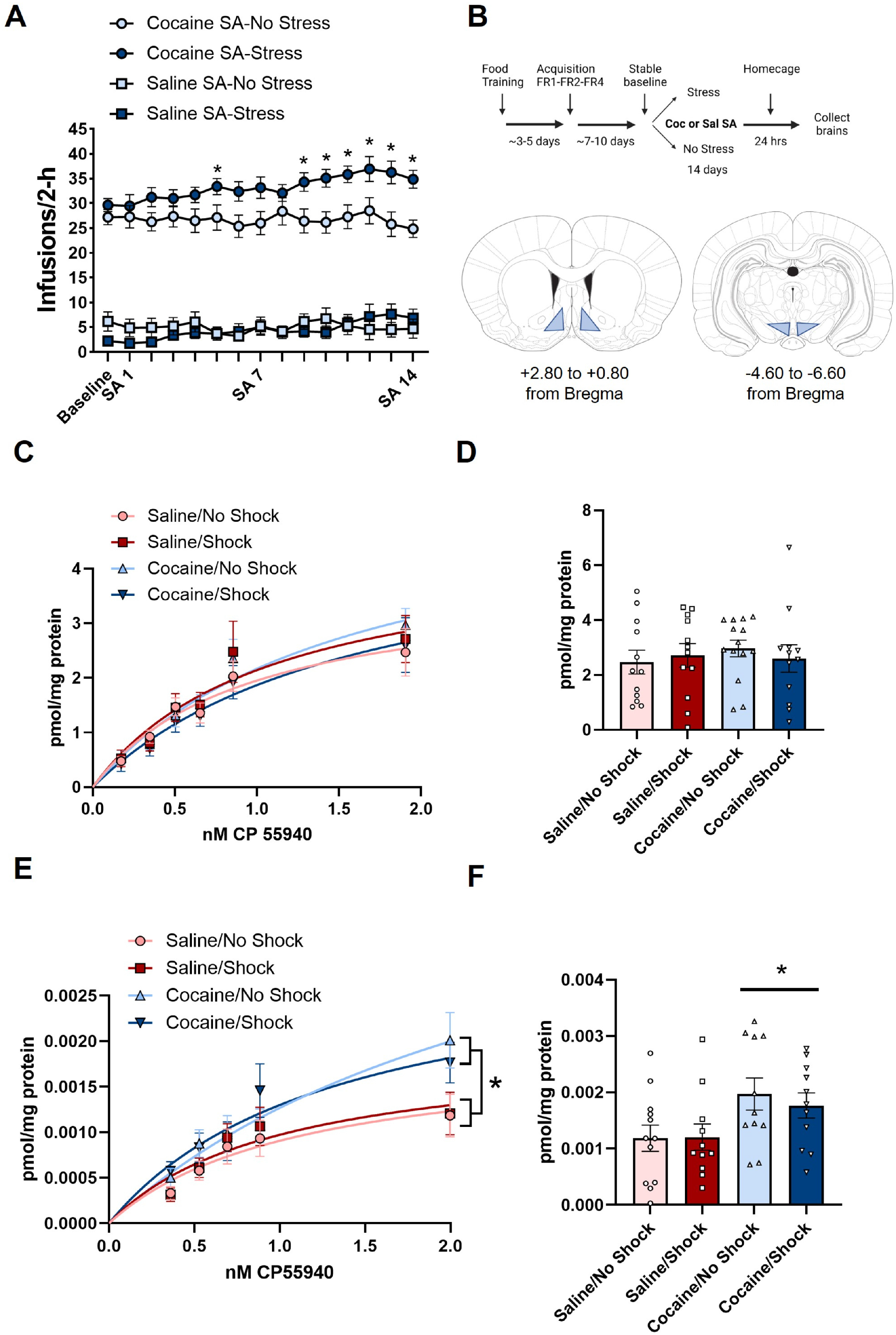
Cocaine self-administration increases cannabinoid receptor type 1 binding in the ventral tegmental area but not nucleus accumbens shell. A) Self-administration data for all animals that were used for the CB1R binding experiment. Repeated daily footshock stress induces a significant escalation of cocaine intake (No Shock, n=14; Shock, n=13) but has no effect on saline self-administration (No Shock, n=14; Shock, n=13; 8p<.05, compared to baseline). B) Experimental timeline and schematic of dissection boundaries for the NAc shell (left) and VTA (right). C) CB1R binding in the NAc shell is similar across all four self-administration conditions as assessed by the full binding curve from the radiolabeled ligand CB1R binding assay. The binding at the highest concentration of CP 55, 940 is represented on the right (D) and there are no differences between groups. E) In the VTA, cocaine self-administration increases CB1R binding. The full binding curves are represented on the left and binding at the highest dose of CP 55,940 is higher in rats that underwent cocaine self-administration (F) but there is no effect of stress on CB1R binding (*p<.05 compared to Saline SA at the 2.0 nM concentration of CP 55,940). Data are presented as mean ± SEM.

### Experiment 5: Cocaine-primed reinstatement following stress-induced escalation of cocaine intake; requirement for CB1R

To determine if stress during cocaine SA produces long-lasting changes that influence reinstatement, rats underwent extinction followed by cocaine-primed reinstatement testing (Fig. 4A). Two-way repeated measures ANOVA revealed an overall effect of extinction day (F_(4,128)_=29.37, p<.001) but no stress x extinction interaction (F_(4,128)_=0.66, p=0.62), indicating no differences in extinction between groups (Fig. 4B). Rats in the shock group took longer to reach extinction criterion (6.00 ± 0.80 days) than controls (4.25 ±0.28 days), but this difference did not reach significance (p=.059). Rats were tested for reinstatement using three cocaine priming doses (2.5, 5, 10 mg/kg, i.p.). Three-way ANOVA revealed a significant cocaine dose x stress x day interaction (F_(3,60)_=6.80, p<.001). While both groups showed significant reinstatement compared to extinction (p<.05, Fig. 4C), rats in the shock group showed greater reinstatement at each cocaine dose tested compared to controls (p<.05).

**Figure 4.**
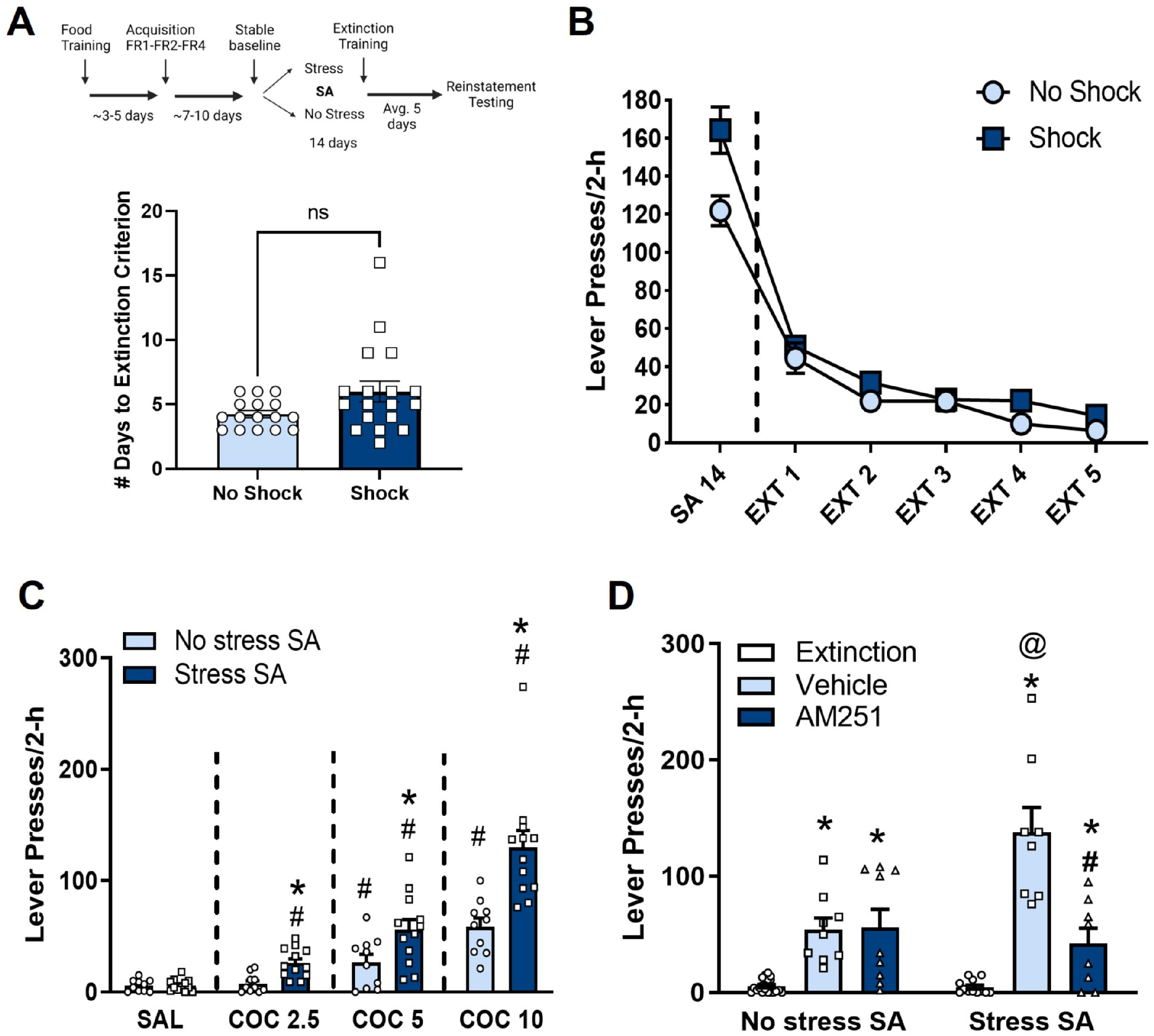
A history of stress-induced escalation of cocaine intake results in increased cocaine-induced reinstatement that is dependent upon CB1R activation. A) Experimental timeline. There is no difference in the number of days to reach extinction criterion between animals with a history of no shock (n=16) or shock (n=18). B) Animals with a history of shock show similar rates of extinction compared to rats with a history of no shock. C) Rats with a history of stress at the time of self-administration (n=12) show increased cocaine-induced reinstatement at multiple doses (2.5, 5, 10 mg/kg, i.p.) compared to rats with a history of no stress at the time of self-administration (n=10; #p<.05, compared to saline; *p<.05, compared to No Shock). D) Systemic administration of AM251 (0, 1 mg/kg, i.p.) 30 min prior to a cocaine injection (10 mg/kg, i.p.) attenuates cocaine-induced reinstatement only in rats with a history of stress at the time of cocaine self-administration (n=8). The same treatment has no effect on cocaine-induced reinstatement in rats with no history of stress at the time of cocaine self-administration (n=9; *p<.05, compared to extinction; @p<.05, compared to No Stress SA-Vehicle; #p<.05, compared to Vehicle). Data are presented as mean ± SEM.

To examine the contribution of CB1R signaling to enhanced reinstatement susceptibility in shocked rats, we assessed the effects of AM251 (1 mg/kg, i.p.) on cocaine-primed reinstatement (10 mg/kg, i.p.). Three-way ANOVA revealed a significant AM251 x stress x day interaction (Fig. 4D; F_(1,15)_=19.64, p<.001). While reinstatement was observed in both groups (p<.001), it was significantly increased in shocked rats (p<.01). Moreover, only shocked rats showed a significant attenuation of cocaine-induced reinstatement by AM251 (p<.001).

## Discussion

### Repeated stress induces an escalation of cocaine self-administration

We previously reported that chronic footshock at the time of daily cocaine SA escalates drug intake but not food-reinforced responding in male rats [16]. Here we extend these findings by demonstrating that escalated cocaine SA persists after chronic stress is terminated and is associated with heightened cocaine-primed reinstatement, despite insignificant differences in extinction compared to non-shocked controls. Thus, our data suggest that the combination of stress and cocaine SA produces unique and lasting neuroadaptations that promote drug use and likely exacerbate SUD severity.

### Role of nucleus accumbens shell and ventral tegmental area CB1R signaling in the regulation of stress-induced escalation of cocaine self-administration

Stress-escalated cocaine SA involves CB1R signaling. Post-stress cocaine SA is selectively reduced by the CB1R antagonist/inverse agonist, AM251, thus restoring SA to levels observed in non-shocked rats. This observation is consistent with findings that another CB1R antagonist, rimonabant, produces pronounced reductions in breakpoints for cocaine when tested under a progressive ratio schedule only in rats that display escalated intake following repeated daily long-access SA [31]. By contrast, CB1R antagonism has minimal effects on cocaine SA under conditions of stable/non-escalated intake [28; 29; 31; 45], although it has been reported that SA is attenuated in CB1R knockout mice [46] but see [47; 48].

Brain endocannabinoid/CB1R signaling is activated by and plays a critical role in adaptation to stress. In addition to buffering the physiological stress response, endocannabinoids promote active coping strategies. In the context of SUDs, such strategies may manifest as heightened drug seeking behavior. Indeed, CB1Rs are expressed throughout the mesocorticolimbic system, including the VTA and NAc shell – regions implicated in stress adaptation and drug SA/seeking [49-52] and key sites for CB1R-dependent reward. Self-administration of THC directly into these regions is prevented by CB1R antagonism, and site-specific THC micro-infusions into these regions produces CB1R-dependent conditioned place preference [53].

Here we provide evidence that CB1R signaling in the NAc shell and VTA contributes to escalated use patterns in rats with a history of footshock stress at the time of cocaine SA. Intra-VTA or intra-NAc shell AM251 micro-infusions selectively attenuated SA in rats that received shock during SA and restored SA to levels observed in non-shocked controls. By contrast, intra-NAc shell AM251 failed to alter SA in non-shocked rats, consistent with findings that intra-NAc shell AM251 reduces SA in rats displaying escalated intake following repeated long-access to cocaine (6-h daily) without affecting SA in short-access rats (1-h daily) [31]. Intra-VTA AM251 did have modest effects on SA in control rats at the highest dose tested; which is consistent with reports that VTA CB1R antagonism attenuates cocaine-induced CPP in mice [54; 55] and cocaine-enhanced motivation assessed in rats self-administering food pellets under a progressive ratio schedule of reinforcement [56]. Additionally, in the VTA, cocaine can mobilize endocannabinoids [43] and cocaine SA upregulates CB1R binding which may also explain why we observe an effect of the high dose of AM251 in the no shock control rats at the high dose of the antagonist.

To determine how the CB1 receptor is altered in rats displaying stress-escalated SA, we assessed CB1R binding parameters in membranes prepared from dissected VTA and NAc shell [57-60]. Neither cocaine SA alone, repeated footshock alone, nor the combination, altered NAc shell CB1R binding. However, cocaine SA increased VTA CB1R binding assessed using a single point analysis regardless of whether rats previously received shock. These findings were unexpected, as previous work found increased CB1R binding in the NAc shell but not VTA following cocaine SA [61]. Considering the large differences in SA observed between shocked rats, which displayed robust escalation, and non-shocked controls, which did not escalate at all, the comparable increase in VTA CB1R binding in both groups was surprising. However, both regions are quite heterogenous and CB1Rs are expressed on multiple cell types. Therefore, cell type-or subregion-specific CB1R expression may be differentially altered in the two groups in a way that better aligns with our behavioral data, but was not detectable with our current dissection strategy and assay. Additionally, future assessment of the roles of endocannabinoid mobilization and downstream CB1R signaling in stress-induced escalation of cocaine intake is warranted.

Based on the established role for dopamine in cocaine SA, we hypothesize that stress-escalated SA potentiates CB1R-mediated activation of dopaminergic signaling via effects in both the VTA and NAc shell. In support of this notion, cocaine mobilizes endocannabinoids in the VTA [43] and NAc shell [29] and CB1R agonists activate mesolimbic neurons [62] and promote NAc dopamine release [63-65]. Moreover, CB1R antagonism attenuates the NAc dopamine response to cocaine [38] and the dopamine response to ip cocaine is blunted in CB1R knockout mice [66].

Given that CB1R signaling is inhibitory to neurotransmitter release, we hypothesize that stress-escalated SA increases activation of CB1R on GABAergic terminals, resulting in disinhibition of mesolimbic neurons. CB1R activation can reduce both inhibitory [67] and excitatory [36] drive onto VTA dopamine neurons. Notably, cocaine exposure promotes CB1R-dependent long-term depression at inhibitory synapses (iLTD) [55] while suppressing CB1R-mediated LTD at excitatory synapses [68] in the VTA, changes consistent with enhanced excitatory drive onto mesolimbic dopamine neurons. Likewise, in the NAc, CB1R activation attenuates both inhibitory [69] and excitatory [70] synaptic transmission and mediates LTD at excitatory synapses [71], including those on parvalbumin+ GABAergic interneurons [72]. As in the VTA, cocaine has been reported to reduce CB1R-dependent LTD in the NAc [73]. While we hypothesize that CB1R signaling promotes escalation and reinstatement through enhanced attenuation of inhibitory transmission in the VTA and NAc, it is possible that alterations in CB1R regulation of excitatory synaptic transmission are also involved.

### CB1R involvement in enhanced cocaine-seeking behavior following stress-induced escalation of cocaine self-administration

Despite no differences in extinction responding between groups, rats that received shock during SA displayed heightened cocaine-primed reinstatement compared to non-shocked controls. These findings are consistent with reports of persistent increases in cocaine-primed reinstatement following stress in rats [74-78], although none of those studies applied stress during a period of ongoing cocaine SA. Prior reports have suggested that the magnitude of drug-primed reinstatement increases according to the amount of previously self-administered cocaine [15; 79; 80]. However, regression analyses failed to demonstrate a relationship between the magnitude of reinstatement and various indicators of earlier SA, such as cumulative intake and area under the curve, suggesting that heightened reinstatement was not solely attributable to increased intake in shocked rats (Fig. S2).

As was the case with SA, cocaine-primed reinstatement was reduced by AM251, but only in rats that received shock during earlier SA. Moreover, AM251 restored cocaine-primed reinstatement to levels observed in controls, suggesting that the heightened susceptibility to drug-primed cocaine seeking was the result of persistent recruitment of endocannabinoid signaling. Prior studies examining the effects of CB1R antagonism on cocaine-primed reinstatement produced mixed results. De Vries et al [28] (iv rimonabant), Xi et al [81] (ip AM251), and Adamczyk et al [45] (ip AM251) reported that CB1R antagonism attenuates cocaine-primed reinstatement, while we [82] (ip AM251) and Kupferschmidt [83] (icv AM251) found no effect on cocaine-primed reinstatement in rats and Ward et al [84] (ip rimonabant) found no effects in mice. Reasons for the discrepancies among rat studies are unclear but may include differences in rat strains/vendors, housing conditions, experimental designs, and treatment parameters. Nonetheless, the findings suggest that the impact of stress during SA on CB1R signaling is long-lasting and may influence relapse susceptibility.

We previously reported that shock-escalated cocaine SA requires elevated glucocorticoids [16]. Moreover, increases in glucocorticoids at the time of SA contribute to both escalated intake and long-term heightened susceptibility to cocaine-primed and stressor-induced reinstatement [17; 18]. Endocannabinoids/CB1R signaling mediate many of the neurobiological and physiological effects of glucocorticoids [56] and our previous studies implicated CB1R/endocannabinoid signaling in the acute effects of corticosterone on cocaine seeking [24]. Thus, we hypothesize that combination of stress and cocaine produces additive or synergistic increases in CB1R activation, which contributes to an escalation of cocaine SA.

### Procedural and experimental design considerations

There are several procedural/design considerations. First, shock-escalated SA persisted for at least five days after the shock was removed from the sessions, which, along with the observation of heightened reinstatement, provides evidence of long-lasting neuroadaptations involving CB1R signaling that promote cocaine seeking. Second, we previously reported that shock failed to escalate food-reinforced behavior (45 mg sucrose pellet [16]), indicating some specificity for cocaine SA. Third, we previously found that the effects of shock require delivery within the drug context and/or at the time of SA; shock escalation was not observed when shock was applied within the SA chambers 4 h after completion of the daily SA sessions or in a different chamber [16]. Fourth, rates of shock-induced escalation varied slightly across cohorts. In some cohorts, escalation was observed within the first session, while in others, it took several days to emerge. The reason for this is unclear. Fifth, escalated SA was observed in each of the four 30-min components during the session and was not confined to the first component of each session when “drug loading” typically occurs. Finally, one limitation is that the present study was conducted using only male rats. Prior work has demonstrated that footshock potentiates the reinstatement of cocaine-induced reinstatement following SA in male but not female rats [85]. Sex differences in endocannabinoid signaling have been reported [86; 87] and extend to endocannabinoid-related effects of chronic stress [88] and cocaine [89]. Considering reports that stress-reactivity differs between cocaine-dependent men and women [90], future studies examining sex differences in CB1R-dependent stress-induced escalation of SA are needed.

### CB1R signaling as a potential therapeutic target for cocaine use disorder

These findings add to the literature demonstrating that adverse life events influence the severity of SUDs. While we found that chronic stress during an ongoing period of SA escalated intake, Miczek and colleagues have demonstrated that a prior history of social defeat intensifies later cocaine SA as observed by more prolonged binge-like behavior when rats are provided unlimited daily access to cocaine [91; 92]. Along with earlier findings [31] these results suggest that the recruitment of mesolimbic endocannabinoid signaling promotes escalating patterns of cocaine use and that heightened CB1R signaling may contribute to CUD. The latter assertion is supported by studies demonstrating that polymorphisms related to the *CN1R* gene, are associated with CUD [93-96] but see [97]. Moreover, the findings indicate that CB1R-targeting medications (e.g., negative allosteric modulators [98; 99] are promising interventions for CUD, particularly for those whose cocaine use is stress related.

## Acknowledgements

We thank Luke Urbanik for assistance with surgeries. Figures were created with BioRender.com

## Author Contributions

JRMc, CJH, and JRM designed the project and experiments. JRMc and JRM supervised all of the experiments. JRMc, CPW, and DMS conducted the surgeries and JRMc ran the behavioral experiments with assistance from CPW, DMS, RS, and JCM. CPW and RS completed the histology. JRMc ran the behavior and collected tissue for the CB1R binding assay and CJH supervised and LAK completed the binding assays and analysis of the binding data. JRMc analyzed and graphed all data. JRMc and JRM wrote the first draft of the manuscript with editing and reviewing by CJH. Funding was acquired by JRMc, CJH, and JRM.

