## Supplemental Information for "Role of mesolimbic cannabinoid receptor 1 in stress-driven increases in cocaine self-administration in male rats"

**Methods**

_Surgery_: For all experiments, rats received indwelling intra-venous catheters for drug self-administration. Each polyurethane catheter (0.6 mm i.d. x 1.1 mm o.d.; Access Technologies, Skokie, IL) was connected to a back-mounted cannula (Plastics One, Roanoke, VA) attached to polypropylene mesh (500 microns; Small Parts, Logansport, IN). Rats were anesthetized with either ketamine HCl (100 mg/kg, i.p.; Henry Schein, Melville, NY) and xylazine (2 mg/kg, i.p.; Henry Schein, Melville, NY) or Isoflurane (2% in $O_2$) and the catheter was implanted into the superior vena cava. The back-mounted cannula was situated approximately 2.5 cm behind the rat's scapula. For rats that received intra-nucleus accumbens (NAc) shell and intra-ventral tegmental area (VTA) infusions, guide cannulas were implanted immediately following the catheter surgery. For intra-nucleus accumbens shell surgeries, the skull was positioned in a stereotaxic frame (Kopf Instruments, Tujunga, CA) and a double-barrel guide cannula apparatus containing two 11 mm stainless steel cannula was implanted 0.5 mm above the NAc shell [0° arm angle; coordinates (in mm): anteroposterior (A/P): + 1.00 from Bregma; mediolateral (M/L): ± 0.75 from midline; dorsoventral (D/V): - 6.50 from the skull surface; incisor bar: -3.3 from interaural line (Paxinos & Watson, 2005)]. For intra-ventral tegmental area surgeries, two single 11 mm stainless steel guide cannula were implanted 0.5 mm above the VTA [12° arm angle; coordinates (in mm): A/P: - 5.40 from Bregma; M/L: ± 2.00 from midline; D/V: - 7.40 from the skull surface; incisor bar: -3.3 from interaural line (Paxinos & Watson, 2005)]. Four anchoring screws were implanted into the skull and both the anchoring screws and guide cannulas were fixed in place with acrylic dental cement. Internal dummy cannulas were inserted to maintain patency. Rats were given Bio-Serv Rimadyl tablets (5 g; Fisher Scientific; Hampton, NH) in their cage for 3 days and Cefazolin antibiotic treatment (100 mg/kg, iv; Henry Schein, Melville, NY) for at least 5 days following surgery. All rats recovered for a minimum of one week before the initiation of self-administration.

_Cocaine self-administration_: All experiments were completed in computer-interfaced operant conditioning chambers. Each chamber was equipped with retractable levers and a stimulus light above each lever in sound-attenuating cubicles and were equipped for delivery of shock through stainless steel grid floors (MED-Associates; Fairfax, VT). Rats were initially food deprived to 90% of their starting weight and were trained to press a lever to receive a sucrose pellet on a fixed ratio (FR) 1 schedule of reinforcement. Once rats acquired lever pressing behavior, they were transitioned to receive an infusion of cocaine (0.5 mg/kg/0.2 mL infusion over 5 s) in response to a lever press on an FR1 schedule during daily self-administration sessions using a modified program (Fig 1A) at which point they were once again given *ad libitum* access to food. The program consisted of four 30-min self-administration blocks wherein the house light was off and the light above the active lever was illuminated. Upon initiation of an infusion of cocaine the light above the lever was extinguished for a 10-s time out period during which lever presses were recorded but not reinforced. Upon conclusion of the time-out period the light above the lever was re-illuminated and lever presses were reinforced once again. The total amount of time that rats were allowed to self-administer cocaine was 2 hrs. Separating the four 30-min self-administration blocks were four 5-min drug-free periods during which the houselight was illuminated, the levers were retracted, and the light above the lever was off. During the self-administration session, a second, inactive lever was extended on which lever presses were recorded but not reinforced. Once rats showed regular responding on an FR1 schedule, they were gradually moved to an FR4 schedule. Stable baseline cocaine intake was established when infusions varied by <10% across 3 days. Following establishment of stable baseline responding, the control no shock group continued to self-administer cocaine under these same parameters for 14 days. The control rats do not receive footshock stress during the drug-free periods.

*Footshock*: The footshock administration consisted of triplets of shock (3 x 0.4 mA; 500 ms duration; 1 s intershock interval) that were given at random intervals during the 5-min period (30 s average inter-triplet shock interval; range 15-60 s). Rats received an average of 7 triplet shocks per 5 min drug-free period for a total average of 28 triplet shocks for the whole self-administration session. The footshock was only administered during the drug-free period and was never administered when drug was freely available.

Effects of CB1R antagonism on cocaine-taking behavior

For experiments in which AM251 was administered, each rat received both the vehicle and AM251. Experiments 2 & 3, rats were tested under “no shock” conditions to isolate the contribution of recruited CB1R signaling to cocaine self-administration and eliminate effects on the acute response to shock. Rats received both the vehicle and AM251 tests in a counter-balanced order with at least one day in between tests and each rat underwent two tests total. Infusions and lever presses were recorded across all four self-administration sessions on the test day.

*Drugs*: Cocaine HCl was obtained from the National Institute of Drug Abuse (NIDA) Drug Supply Program. Cocaine was dissolved in bacteriostatic saline (0.9%). AM251 (Sigma-Aldrich, St. Louis, MO), when administered systemically, was first dissolved in ethanol, followed by Cremaphor and finally by basteriostatic saline (0.9%) in a 1:1:18 ratio. AM251, when administered intra-cranially, was dissolved in 100% DMSO.

*Intra-cranial drug administration*: The CB1R antagonist/inverse agonist AM251 (0, 1, 3 µg/side) was micro-infused bilaterally directly into the NAc shell or VTA 15-min prior to the start of the self-administration as described in Experiment 4. For the NAc shell, infusion needles were comprised of a dual injector containing two 11.5 mm 30-gauge stainless steel injectors (Plastics One, Roanoke, VA) attached to polyethylene (PE)-20 tubing and Hamilton syringes. For the VTA, infusion needles were comprised of single 11.5 mm 30-gauge stainless steel injectors attached to PE-20 tubing and Hamilton syringes. For both brain regions, the injectors extended 0.5 mm beyond the guide cannula. AM251 or vehicle was backfilled into the infusion needle and all drugs were infused at a volume of 0.3 µL/side and at a rate of 0.3 µL/min using a syringe pump. The needles were kept in place for an additional 1-min to allow diffusion. Internal dummy injectors were placed back into the guide cannula following the infusion to maintain cannula patency in between tests.

*Histology*: To determine guide cannula placement, rats were euthanized with $CO_2$ and brains were removed and post-fixed in 4% paraformaldehyde for at least 2 weeks. Brains were then cryoprotected in 30% sucrose and 50-µm sections were taken at the level of the NAc shell and the VTA using a cryostat and stored in PBS. Tissue sections were mounted onto slides with a 0.3% gelatin solution and were stained using a cresyl violet nuclear stain. Guide cannula placement was determined to be correct if the injector terminated in the NAc shell or VTA in both hemispheres. Any rat that did not have correct bilateral placement or that contained extensive damage at the injection site were excluded from study (NAc shell, n=9; VTA, n=6).

*CB1R binding assay*:

Tissue collection: Rats underwent self-administration under “no stress” or “stress” conditions for 14 days, as described in Experiment 1. To examine potential changes in CB1R binding as a consequence of either cocaine self-administration or the repeated stressor alone, additional control groups were run. These control groups were exposed to similar testing conditions as described in Experiment 1 but were provide access to intravenous saline instead of cocaine. Briefly, following food training, rats were allowed to self-administer a

0.9% saline solution on an FR1 schedule. “Saline SA” rats were given 7-8 days of additional access to match the acquisition period for the cocaine SA rats. These rats were then allowed to self-administer saline on an FR1 schedule for 14 days under no stress or stress conditions. 24-h after the last self-administration session, rats were rapidly sedated with isoflurane for approximately 30-s. The rats were decapitated and brains were removed and rapidly flash frozen in 2-methylbutane chilled on dry ice (<2-min from decapitation to frozen). Brains were then stored at -80°C until processing. To dissect the NAc shell and VTA, 2-mm thick frozen sections were taken using a cold metal brain block on dry ice. For the NAc shell dissections, a 2-mm thick section was cut from A/P +2.80 mm to +0.80 mm from Bregma and a scalpel was used to dissect the nucleus accumbens shell medial to the nucleus accumbens core. For the VTA dissections, a 2-mm thick section was cut from A/P -4.60 mm to -6.60 mm from Bregma and a scalpel was used to dissect the VTA. The dissected tissue was collected in microcentrifuge tubes and stored at -80°C until processing.

Membrane Preparation

Dissected brain sections were homogenized in 10 volumes of TME buffer (50 mM Tris Base, 1 mM EDTA, 2 mM $MgCl_2$, pH 7.4). The homogenates were centrifuged at 12,000 rpm for 20 min at 4°C after which the supernatant was rapidly decanted. The remaining pellet, the membrane fraction, was resuspended in 5 µL/mg starting weight TME buffer. Protein concentrations were determined by the Bradford method.

CB1R binding assay:

CB1R receptor binding assays were performed using a Multi-screen Filtration System with Durapore 1.2-µM filters (Millipore, Bedford, MA) as described previously (Hillard et al, 1995). Incubations (total volume = 200 µL) were carried out using TME buffer containing 1 mg/mL bovine serum albumin (BSA). Membranes (10 µg protein per incubate) were added to the wells containing 0.25, 0.5, 0.75, 1.0, 1.25, or 2.0 nM $^3$H-CP 55,940, a CB1R agonist. AM251 (12 µM) was used to determine nonspecific binding. $K_D$ and $B_{max}$ values for each group were determined by nonlinear curve fitting to the single site binding equation using GraphPad Prism (San Diego, CA). We initially analyzed binding curves for each group with one-site specific binding using least squares fit nonlinear regression. We then compared fits using the extra sum-of-squares approach which identified whether one curve could adequately fit the data from all groups.

Extinction/Reinstatement

Following 14 days of self-administration under “stress” or “no stress” conditions, rats underwent extinction training, during which cocaine solution was replaced with saline and responding on the active lever gradually decreased. Rats underwent extinction training until the extinction criterion was met (<15 lever presses/2 hr). Once rats met this criterion, a reinstatement test was conducted the following day. On the day of the reinstatement test, rats were given a systemic injection of cocaine (2.5, 5, 10 mg/kg, i.p.) or saline immediately prior to the beginning of the reinstatement session. A complete within-subjects design was used wherein all rats underwent multiple reinstatement tests in a counterbalanced fashion for a maximum of 4 reinstatement tests. Rats that did not receive all doses of cocaine were excluded from the study. Rats were given additional extinction/washout sessions between doses and were required to reach extinction criterion again before an additional reinstatement test was administered. For the AM251 experiments, on the reinstatement test day, additional groups of rats first received an injection of AM251 (1 mg/kg, ip) or vehicle followed 30 min later with an injection of cocaine (10 mg/kg, i.p.) immediately prior to the start of the reinstatement test session. As above, rats received additional extinction training days in between reinstatement tests to ensure that they reached the extinction criteria (<15 lever presses/2-h session) prior to undergoing an additional reinstatement test. Each rat received no more than 3 tests total.

*Statistical analysis*: Statistical analyses were conducted using SPSS (IBM Analytics, Armonk, NY) or GraphPad Prism (San Diego, CA). For behavioral experiments, cocaine infusions or lever presses were analyzed using two- or three-way repeated measures ANOVA followed by Holm-Sidak or Dunnett’s post hoc testing where appropriate. For CB1R binding experiments, binding at the highest concentration of CP 55,940 was analyzed using a 2-way repeated measures ANOVA.

**Results**

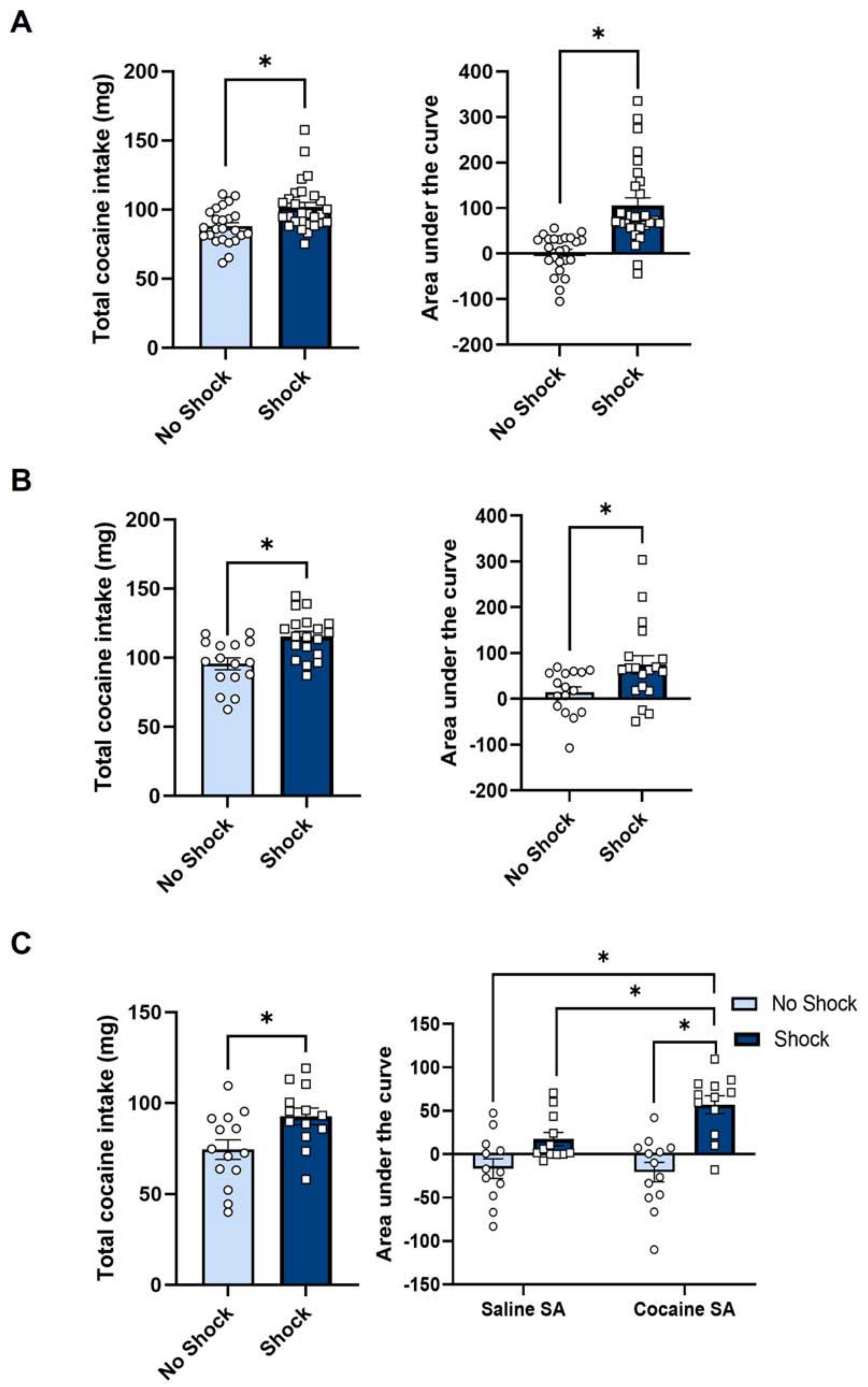
A
Total cocaine intake (mg)
Area under the curve
No Shock
Shock
*
B
Total cocaine intake (mg)
Area under the curve
No Shock
Shock
*
C
Total cocaine intake (mg)
Area under the curve
No Shock
Shock
Saline SA
Cocaine SA
*

**Figure S1**. Total cocaine intake and area under the curve for self-administration in Figures 1-3. Repeated shock stress at the time of cocaine self-administration increases total cocaine intake compared to no shock animals from self-administration that is represented in Figure 1 (A), Figure 2 (B), and Figure 3 (C). Area under the curve (AUC) was measured for self-administration from days 1-14 with baseline self-administration set as 0 and shock animals have greater AUC compared to no shock animals for self-administration that is presented in Figure 1 (A) and Figure 2 (B). Repeated shock stress administered at the time of cocaine self-administration resulted in increased AUC for self-administration represented in Figure 3 (C) compared to cocaine SA-no shock, saline SA-no shock, and saline SA-shock (*p<.05). Data are presented as mean ± SEM.

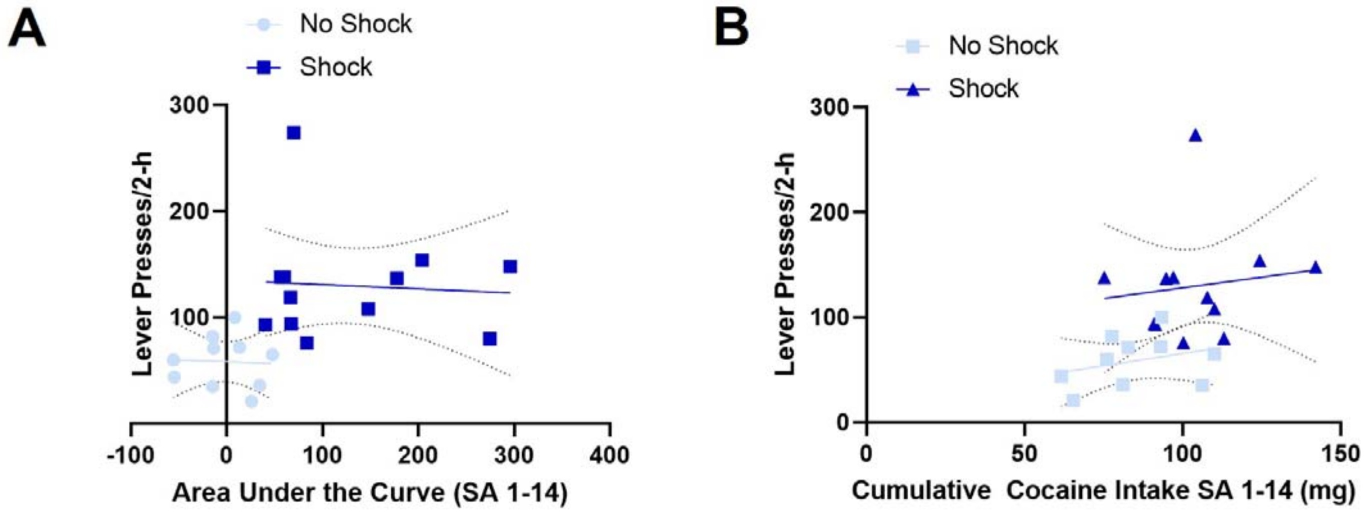


**Figure S2**. Linear regression test of whether measures of self-administration in the shock and no shock groups predicted reinstatement to a high dose of cocaine (10 mg/kg, i.p). A) Simple linear regression was used to test if area under the curve (SA 1-14) of infusions across SA significantly predicted lever presses during the cocaine-primed reinstatement test. The overall regression was not statistically significant for either the no shock ($R^2$=0.002, $F_{(1,8)}$=0.018, p=0.89) or the shock group ($R^2$=0.004, $F_{(1, 10)}$ = 0.046, p=0.83). B) Simple linear regression was used to test if cumulative cocaine intake across SA days 1-14 (in mg) significantly predicted lever presses during the cocaine-primed reinstatement test. The overall regression was not statistically significant for either the no shock ($R^2$=0.093, $F_{(1,8)}$=0.826, p=0.39) or the shock group ($R^2$=0.017, $F_{(1, 10)}$ = 0.182, p=0.68).

**Tables**

| ***Comparison*** | ***Statistical Test*** | ***Statistic*** | ***p-value*** |
|---|---|---|---|
| **B** | | | |
| **Infusions across SA days** | Two-way RM ANOVA | | |
| Stress (between) x SA Day (within) | Interaction | F(14,700)=4.72 | **P<.0001** |
| Day | Main effect | F(14,700)=3.47 | **P<.0001** |
| Stress: No shock, n=24; Shock, n=28 | Main effect | F(1,50)=9.30 | **P<.005** |
| No Shock: Baseline:SA 1-14 | Dunnett's post-hoc test | | P>0.9 |
| Shock: Baseline: SA 1-14 | Dunnett's post-hoc test | | **P<.01** |
| **C** | | | |
| **Active lever presses across SA days** | Two-way RM ANOVA | | |
| Stress (between) x SA Day (within) | Interaction | F(14,700)=2.56 | **P<.005** |
| Day | Main effect | F(14,700)=1.86 | **P<.05** |
| Stress: No shock, n=24; Shock, n=28 | Main effect | F(1,50)=7.61 | **P<.01** |
| No Shock: Baseline:SA 1-14 | Dunnett's post-hoc test | | P>0.9 |
| Shock: Baseline: SA 1-14 | Dunnett's post-hoc test | | **P<.01** |
| **Inactive lever presses across SA days** | Two-way RM ANOVA | | |
| Stress (between) x SA Day (within) | Interaction | F(14,700)=1.54 | P=0.09 |
| Day | Main effect | F(14,700)=1.14 | P=0.32 |
| Stress: No shock, n=24; Shock, n=28 | Main effect | F(1,50)=0.47 | P=0.49 |
| **D** | | | |
| **Intake across SA block across SA days: No Shock, n=24** | Two-way RM ANOVA | | |
| SA Block (within) x SA Day (within) | Interaction | F (42, 966) = 1.141 | P=0.25 |
| SA Block | Main effect | F (3, 69) = 0.2569 | P=0.86 |
| SA Day | Main effect | F (14, 322) = 0.9796 | P=0.47 |
| **E** | | | |
| **Intake across SA block across SA days: Shock, n=28** | Two-way RM ANOVA | | |
| SA Block (within) x SA Day (within) | Interaction | F (42, 1134) = 1.343 | P=0.07 |
| SA Block | Main effect | F (3, 81) = 0.09143 | P=0.96 |
| SA Day | Main effect | F (14, 378) = 5.909 | **P<0.0001** |
| **F** | | | |
| **Intake across SA days: SA 14-P5** | | | |
| Stress (between) x SA Day (within) | Interaction | F(5,130)=0.22 | P=0.95 |
| Day | Main effect | F(5,130)=0.37 | P=0.87 |
| Stress; No Shock=13; Shock=15 | Main effect | F(1,26)=7.43 | **P<0.01** |
| **G** | | | |
| **Effect of AM251 on infusions** | Two-way RM ANOVA | | |
| Drug (Within) x Stress (Between) | Interaction | F(1,14)=18.56 | **P<0.001** |
| Drug (Vehicle vs AM251) | Main effect | F (1, 14) = 8.667 | **P<0.01** |
| Stress (No Stress vs Stress) | Main effect | F (1, 14) = 0.2680 | P=0.61 |
| No Shock: Vehicle-AM251; n=9 | Holm-Sidak post hoc test | | P=0.32 |
| Shock: Vehicle-AM251; n=7 | Holm-Sidak post hoc test | | **P<0.01** |

**Table S1**. Detailed statistics for Figure 1. Significant P-values are bolded.

| ***Comparison*** | ***Statistical Test*** | ***Statistic*** | ***p-value*** |
|---|---|---|---|
| **B** | | | |
| **Infusions across SA days** | Two-way RM ANOVA | | |
| Stress (between) x SA Day (within) | Interaction | F (14, 462) = 0.741 | P=0.73 |
| Day | Main effect | F (14, 462) = 1.423 | P=0.14 |
| Stress: No shock, n=16; Shock, n=19 | Main effect | F (1, 33) = 12.83 | **P<0.001** |
| No Shock: Baseline:SA 1-14 | Dunnett's post-hoc test | | P>0.9 |
| Shock: Baseline: SA 3, 5, 6, 7, 12, 14 | Dunnett's post-hoc test | | **P<0.05** |
| **C** | | | |
| **Effect of intra-NAc shell AM251 on infusions** | Three-way RM ANOVA | | |
| Drug (Within) x Stress (Between) x Dose (Between) | Interaction | F(1,28)=7.59 | **P<0.05** |
| Drug x Stress | Interaction | F(1,28)=45.37 | **P<0.001** |
| Drug x Dose | Interaction | F(1,28)=7.59 | **P<0.01** |
| Stress x Dose | Interaction | F(1,28)=0.56 | P=0.46 |
| Drug (Vehicle vs AM251) | Main effect | F (1, 28) = 55.22 | **P<0.001** |
| Stress (No Stress vs Stress) | Main effect | F (1, 28) = 6.86 | **P<0.05** |
| Dose (1µg vs 3 µg) | Main effect | F(1,28)=0.38 | P=0.46 |
| Vehicle: No Shock (n=16)-Shock (n=16) | Holm-Sidak post hoc test | | **P<0.001** |
| No Shock: Vehicle-AM251 (1µg); n=9 | Holm-Sidak post hoc test | | P=0.69 |
| No Shock: Vehicle-AM251 (3µg); n=7 | Holm-Sidak post hoc test | | P=0.69 |
| Shock: Vehicle-AM251 (1µg); n=8 | Holm-Sidak post hoc test | | **P<0.01** |
| Shock: Vehicle-AM251 (3µg); n=8 | Holm-Sidak post hoc test | | **P<0.01** |
| **E** | | | |
| **Effect of intra-VTA AM251 on infusions** | Three-way RM ANOVA | | |
| Drug (Within) x Stress (Between) x Dose (Between) | Interaction | F(1,23)=0.04 | P=0.84 |
| Drug x Stress | Interaction | F(1,23)=4.31 | **P<0.05** |
| Drug x Dose | Interaction | F(1,23)=0.31 | P=0.59 |
| Stress x Dose | Interaction | F(1,23)=0.39 | P=0.54 |
| Drug (Vehicle vs AM251) | Main effect | F (1, 23) = 22.12 | **P<0.001** |
| Stress (No Stress vs Stress) | Main effect | F (1, 23) = 3.74 | **P<0.05** |
| Dose (1µg vs 3 µg) | Main effect | F(1,23)=0.73 | P=0.40 |
| Vehicle: No Shock (n=13)-Shock (n=14) | Holm-Sidak post hoc test | | **P<0.01** |
| No Shock: Vehicle-AM251 (1µg); n=6 | Holm-Sidak post hoc test | | P=0.19 |
| No Shock: Vehicle-AM251 (3µg); n=7 | Holm-Sidak post hoc test | | **P<0.05** |
| Shock: Vehicle-AM251 (1µg); n=6 | Holm-Sidak post hoc test | | **P<0.05** |
| Shock: Vehicle-AM251 (3µg); n=8 | Holm-Sidak post hoc test | | **P<0.05** |

**Table S2**. Detailed statistics for Figure 2. Significant P-values are bolded.

| ***Comparison*** | ***Statistical Test*** | ***Statistic*** | ***p-value*** |
|---|---|---|---|
| **B** | | | |
| **Infusions across SA days** | Two-way RM ANOVA | | |
| SA Condition (between) x SA Day (within) | Interaction | F (42, 700) = 1.58 | **P<.05** |
| Day | Main effect | F (14, 700) = 2.82 | **P<0.001** |
| SA Condition: Coc/No shock, n=14; Coc/Shock, n=13; Sal/No Shock, n=14; Sal/Shock, n=13 | Main effect | F (3, 50) = 98.77 | **P<0.001** |
| Coc/No Shock: Baseline:SA 1-14 | Dunnett's post-hoc test | | P>0.9 |
| Coc/Shock: Baseline: SA 11, 12, 13, 14 | Dunnett's post-hoc test | | **P<0.05** |
| Sal/No Shock: Baseline: SA 1-14 | Dunnett's post-hoc test | | P>0.50 |
| Shock: Baseline: SA 13, 14 | Dunnett's post-hoc test | | **P<0.05** |
| **D** | | | |
| **Single point binding - NAc Shell** | Two-way ANOVA | | |
| Stress (Between) x SA Condition (Between) | Interaction | F(1,46)=0.54 | P=0.47 |
| SA Condition (Coc SA vs Sal SA) | Main effect | F (1, 46) = 0.22 | P=0.64 |
| Stress (No Stress vs Stress) | Main effect | F (1, 46) = 0.02 | P=0.88 |
| **F** | | | |
| **Single point binding - VTA** | Two-way ANOVA | | |
| Stress (Between) x SA Condition (Between) | Interaction | F(1,41)=0.21 | P=0.65 |
| SA Condition (Coc SA vs Sal SA) | Main effect | F (1, 41) = 7.49 | **P<.01** |
| Stress (No Stress vs Stress) | Main effect | F (1, 41) = 0.14 | P=0.71 |

**Table S3**. Detailed statistics for Figure 3. Significant P-values are bolded.

| *Comparison* | *Statistical Test* | *Statistic* | *p-value* |
|---|---|---|---|
| A | | | |
| **# Days to reach extinction criterion** | Unpaired t-test | t(32)=1.96 | P=0.059 |
| B | | | |
| **Active lever presses across extinction days** | Two-way RM ANOVA | | |
| Stress (between) x SA Day (within) | Interaction | F(4,128)=0.66 | P=0.62 |
| Day | Main effect | F(4,128)=29.37 | **P<0.001** |
| Stress: No shock, n=16; Shock, n=18 | Main effect | F(1,32)=2.77 | P=0.11 |
| C | | | |
| **Cocaine-induced (0, 2.5, 5, 10 mg/kg) reinstatement** | Three-way RM ANOVA | | |
| Day (Within) x Coc Dose (Within) x Stress (Between) | Interaction | F (3, 60) = 6.80 | **P<0.001** |
| Coc Dose x Stress | Interaction | F (3, 60)=9.12 | **P<0.001** |
| Day x Stress | Interaction | F(1,20)=15.32 | **P<0.001** |
| Coc Dose x Day | Interaction | F(3,60)=54.64 | **P<0.001** |
| Coc Dose (0, 2.5, 5, 10 mg/kg) | Main Effect | F(3,60)=53.19 | **P<0.001** |
| Day (Extinction vs Reinstatement) | Main effect | F(1,20)=101.31 | **P<0.001** |
| Stress (No Stress: n=10 vs Stress: n=12) | Main effect | F (1,20) = 23.74 | **P<0.001** |
| Coc 2.5: No Stress-Stress | Holm-Sidak | | **P<0.001** |
| Coc 5: No Stress-Stress | Holm-Sidak | | **P<0.05** |
| Coc 10: No Stress-Stress | Holm-Sidak | | **P<0.001** |
| No Stress-Coc 2.5: Ext vs Rst | Holm-Sidak | | P=0.65 |
| No Stress-Coc 5: Ext vs Rst | Holm-Sidak | | **P<0.05** |
| No Stress-Coc 10: Ext vs Rst | Holm-Sidak | | **P<0.01** |
| Stress-Coc 2.5: Ext vs Rst | Holm-Sidak | | **P<0.001** |
| Stress-Coc 5: Ext vs Rst | Holm-Sidak | | **P<0.001** |
| Stress-Coc 10: Ext vs Rst | Holm-Sidak | | **P<0.001** |
| D | | | |
| **Effect of AM251 on Cocaine-induced (10 mg/kg) reinstatement** | Three-way RM ANOVA | | |
| Day (Within) x Drug (Within) x Stress (Between) | Interaction | F(1,15) = 19.64 | **P<0.001** |
| Drug x Stress | Interaction | F(1,15) = 19.95 | **P<0.001** |
| Day x Stress | Interaction | F(1,15) = 3.06 | P=0.10 |
| Drug x Day | Interaction | F(1,15) = 20.59 | **P<0.001** |
| Drug (Vehicle vs AM251) | Main Effect | F(1,15) = 16.12 | **P<0.001** |
| Day (Extinction vs Reinstatement) | Main effect | F(1,15) = 44.34 | **P<0.001** |
| Stress (No Stress: n=9 vs Stress: n=8) | Main effect | F(1,15) = 3.45 | P=0.08 |
| Vehicle: No Shock - Shock | Holm-Sidak | | **P<0.01** |
| No Stress: Veh-AM251 | Holm-Sidak | | P=0.89 |
| Stress: Veh-AM251 | Holm-Sidak | | **P<0.001** |
| No Stress-Vehicle: Ext vs Rst | Holm-Sidak | | **P<0.001** |
| No Stress-AM251: Ext vs Rst | Holm-Sidak | | **P<0.01** |
| Stress-Vehicle: Ext vs Rst | Holm-Sidak | | **P<0.001** |
| Stress-AM251: Ext vs Rst | Holm-Sidak | | **P<0.05** |

**Table S4.** Detailed Statistics for Figure 4. Significant P-values are bolded.

| | | Saline SA/No Shock | Saline SA/Shock | Cocaine SA/No Shock | Cocaine SA/Shock |
|---|---|---|---|---|---|
| $B_{max}$ (pmol/mg protein) | NAc Shell | 3.80 ± 0.79 | 4.47 ± 1.26 | 5.57 ± 1.29 | 4.92 ± 1.69 |
| | VTA | 1.92 ± 0.65 | 1.95 ± 0.61 | 4.72 ± 2.11 | 3.02 ± 0.76 |
| $K_D$ (nM) | NAc Shell | 0.94 ± 0.38 | 1.08 ± 0.58 | 1.57 ± 0.61 | 1.64 ± 0.94 |
| | VTA | 1.12 ± 0.72 | 1.01 ± 0.61 | 2.73 ± 1.78 | 1.33 ± 0.76 |

**Table S5**. $B_{max}$ and $K_D$ values from the CB1R binding assay in the NAc shell or VTA by SA group. Results are presented as mean ± SEM.
